# Selective brain hypothermia ameliorates focal cerebral ischemia-reperfusion injury via inhibiting Fis1 in rats

**DOI:** 10.1101/623611

**Authors:** Yanan Tang, Huailong Chen, Xiaopeng Sun, Weiwei Qin, Fei Shi, Lixin Sun, Xiaona Xu, Gaofeng Zhang, Mingshan Wang

**Affiliations:** Department of Anesthesiology, Affiliated Qingdao Municipal Hospital of Qingdao University, Qingdao 266071, P.R. China; Department of Central Laboratory, Affiliated Qingdao Municipal Hospital of Qingdao University, Qingdao 266071, P.R. China

## Abstract

Mitochondrial immoderate fission induces neuronal apoptosis following focal cerebral ischemia-reperfusion (I/R) injury. With fewer complications, selective brain hypothermia (SBH) is considered an effective treatment against neuronal injury after stroke. However, the specific mechanism by which SBH influences mitochondrial fission remains unknown. Mitochondrial fission 1 protein (Fis1), a key factor of the mitochondrial fission system, regulates mitochondrial dynamics. This study aimed to investigate whether SBH regulates Fis1 expression in focal cerebral I/R injury. In this study, rat middle cerebral artery occlusion (MCAO) models were established. After 2 h of occlusion, cold saline or normal saline was pumped into rats in different groups through the carotid artery, followed by restoration of blood perfusion. Cortical and rectal temperatures showed that the cold saline treatment achieved SBH. Cerebral I/R injury increased neurological deficit scores (NDS); neuronal apoptosis; Fis1 protein and mRNA expression; cytosolic cytochrome c (cyto-Cyto c) protein expression at 6, 24 and 48 h postreperfusion; and cerebral infarct volumes at 24 h postreperfusion. Interestingly, SBH inhibited Fis1 protein and mRNA expression, blocked cyto-Cyto c protein expression, preserved neuronal cell integrity, and reduced neuronal apoptosis. However, normal saline treatment rarely resulted in positive outcomes. Based on these results, SBH inhibited Fis1 expression, thus ameliorating focal cerebral I/R injury in rats.

## Introduction

Stroke is one of the leading causes of death and disability in the world [1–3]. Ischemic stroke caused by cerebral thrombosis or endovascular embolization accounts for a major portion of strokes. Early restoration of blood perfusion is the most effective treatment for stroke; this treatment provides nutrients and oxygen and removes toxic metabolites [4,5]. However, vessel recanalization often exacerbates the existing tissue damage [6,7]; thus, this condition is called cerebral ischemia-reperfusion (I/R) injury. Recent studies have suggested that morphological changes in mitochondria might be relevant to I/R injury [8,9]. Within the cell, mitochondria exist in an ever-changing dynamic state—constant fission and fusion—to form mitochondrial networks and maintain cellular physiological function and survival [10]. Notably, growing evidence has indicated that I/R injury can break this balance to induce neuronal apoptosis [11–13]. Mitochondrial fission protein 1 (Fis1), a 16-kDa protein anchored to the outer membrane of the mitochondria, mediates mitochondrial fission by recruiting cytoplasmic dynamin-related protein1 (Drp1) into the mitochondrial outer membrane [14]. Overexpression of Fis1 increases the frequency of mitochondrial fission, subsequently increasing the release of Cyto c and disrupting mitochondrial membrane potential, thus inducing neuronal apoptosis [15]. In addition, in cells with the Fis1 deletion mutation, Drp1 is mostly localized to the cytoplasm, and mitochondrial fission is inhibited [16]. In conclusion, these studies indicate that Fis1 is a key factor regulating mitochondrial fission.

Hypothermia may be the most potent neuroprotective strategy that can affect multiple pathways at various stages of ischemic stroke, such as calcium overload, oxidative stress, mitochondrial dysfunction, and apoptosis. According to our previous study, general hypothermia-induced neuroprotective effects against cerebral I/R injury are associated with suppressing mitochondrial fission by inhibiting the translocation of Drp1 from the cytoplasm to mitochondria in mice [17]. However, compared with general hypothermia, selective brain hypothermia (SBH) is strongly expected to become a novel attractive treatment for ischemic stroke due to its rapid cooling action and fewer systemic side effects [18,19]. Unfortunately, whether SBH could alleviate I/R injury by inhibiting Fis1 is not well known.

Therefore, in this study, we established a model of focal cerebral I/R using the intraluminal filament technique to embolize the middle cerebral artery (MCA) in rats. Moreover, we induced SBH by infusing cold saline through the internal carotid artery (ICA) in rats before reperfusion. In addition, we explored whether SBH could alleviate cerebral I/R injury by inhibiting Fis1 expression.

## Materials and methods

### Experimental animals and groups

A total of 160 healthy male Sprague-Dawley rats (SPF grade, weighing 200-250 g, aged 8-12 weeks) were provided by the Animal Company of Da Ren Fucheng (Qingdao, Shandong province, China) (license No: SCXKL [LU] 2014-007). All the experimental procedures were performed strictly in accordance with the relevant regulations of the Care and Use of Laboratory Animal from the National Institutes of Health. Experiments were authorized by the ethics committee of Qingdao Municipal Hospital of China (approval No. 2019008). Animals were housed with five animals per cage with free access to water and food. The rats were kept in a room under standard laboratory conditions with a temperature of 18-24 °C, humidity of 50-60%, and a 12-hour light/dark cycle. The rats were randomly divided into 4 groups as follows (n=40 each group): sham group, I/R group, HT group (I/R+cold saline) and NT group (I/R+warm saline).

### Establishment of the focal cerebral I/R injury model

Focal cerebral ischemia was induced by transient MCAO using the intraluminal filament technique as described in S1 Methods. The rectal temperature of the rats was maintained at 36.8-37.2 °C by a heating plate throughout the operation. After blocking the right MCA for 2 h, the filament was slowly pulled out to allow blood reperfusion. In the sham group, the carotid arteries were exposed without obstructing blood flow.

### Establishment of SBH

According to previously used successful parameters [19], 4 °C cold saline was infused (20 ml/kg) through the microcatheter placed in the right ICA via the external carotid artery for 15 min immediately after removal of the filament in the HT group. To eliminate the interference from hemodilution by saline infusion, we performed 37 °C warm saline infusion in the same manner in the NT group. During saline infusion, cortical and rectal temperatures were monitored. Needle thermistor probes (BAT-12 Microprobe Thermometer; Physitemp Instruments, Inc., NJ USA) were placed into the cortex through holes made 3 mm lateral to bregma, 3 mm posterior to bregma, and 3 mm lateral to bregma on the ipsilateral side. Body temperatures were measured through the rectum. Then, rats were returned to their cages with free access to food and water and were closely monitored.

### Evaluation of neurological deficits

When rats’ respiratory and heart rates were stable after reperfusion, we evaluated the neurological deficits of the rats using the Zea Longa 5-point scoring method [20]. Scores ranging from 1 to 3 points were considered an indicator of the successful establishment of the MCAO model; rats with other scores were excluded. The excluded rats were replaced in subsequent experiments. At 6, 24 and 48 h postreperfusion, the rats in each group were evaluated for neurological deficits by the relevant index.

### Hematoxylin-eosin (HE) staining and terminal deoxynucleotidyl transferase-mediated dUTP nick-end labeling (TUNEL) staining

At 6, 24 and 48 h postreperfusion, the rats were euthanized with pentobarbital sodium and transcardially perfused with 0.9% NaCl, followed by 4% paraformaldehyde (PFA). Brains were quickly removed, and brain tissues at the coronal plane from 1 to 4 mm posterior to the optic chiasma were preserved. Then, the brains were soaked in 10% paraformaldehyde phosphate buffer overnight, dehydrated, cleared, dipped in wax, embedded, and cut into 5-μm-thick coronal paraffin sections. HE staining was performed as follows: hematoxylin staining for 10 min, 75% hydrochloric acid alcohol solution for 30 seconds of decoloring, eosin staining for 10 min and 90% ethanol for 35 seconds of decoloring. Six visual fields were randomly selected and observed in the ischemic penumbra of each brain slice under a 400× magnification objective lens (Olympus, Tokyo, Japan). TUNEL staining was used to detect neuronal apoptosis. Briefly, paraffin sections were dewaxed, hydrated, and subjected to TUNEL staining. The specific steps were carried out according to the instructions of a TUNEL kit (Merck Millipore, USA). Five regions were randomly selected from the ischemic penumbra of each brain slice under a 400× magnification objective lens (Olympus, Tokyo, Japan). The mean values were calculated to determine the number of TUNEL-positive cells, which exhibited brownish yellow granules in the nucleus.

### Western blot analysis

At 6, 24 and 48 h postreperfusion, brains were quickly removed, and ischemic penumbra cortices were rapidly dissected on ice as previously reported [21], the specific steps as described in S2 Methods. Mitochondrial and cytosolic fractions were separated using a cytosol/mitochondria fractionation kit (Beyotime Biotechnology, China) according to the manufacturer’s instructions. The cytosol fraction was obtained at 4 °C for Cyto c assays. The expression of specific markers (Cox-IV for mitochondria) was used to ensure the purity of the cytosol fraction. The brain ischemic penumbra cortices were homogenized in lysis buffer (RIPA and PMSF) and then centrifuged at 12000 g for 15 min at 4 °C. The samples were used for Fis1 assays.

The protein concentration was measured with a bicinchoninic acid (BCA) protein assay kit (Beyotime Biotechnology, China). Equal amounts of the protein samples (30~50 μg) were loaded in each well of 10% sodium dodecyl sulfate-polyacrylamide gels, separated by electrophoresis, and transferred onto a polyvinylidene fluoride (PVDF) membrane by the semi-dry method. After washing with 1× TBST and blocking with 3% BSA at room temperature for 2 h, the membranes were incubated overnight at 4 °C with the following primary antibodies: rabbit anti-rat Fis1 monoclonal antibody (1:1000, Abcam, USA), rabbit anti-rat Cyto c monoclonal antibody (1:1000, Cell Signaling Technology, USA), rabbit anti-rat Cox-IV monoclonal antibody (1:2000, Abcam, USA) and rabbit anti-rat β-actin monoclonal antibody (1:5000, Zhongshan Goldenbridge Biotechnology, China). After washing primary antibodies with 1× TBST, the membranes were incubated with goat anti-rabbit HRP-conjugated secondary antibody (1:5000, Zhongshan Goldenbridge Biotechnology, China) in blocking solution for 1 h at room temperature. The membranes were washed again, and the immunoreactive bands were developed using ECL (Beyotime Biotechnology, China). The images were quantified by ImageJ software (NIH, Maryland, USA), and the expression levels of Fis1 and cyto-Cyto c were reflected by the ratios of Fis1 and cyto-Cyto c to the β-actin band gray value, respectively.

### Quantitative reverse transcription-polymerase chain reaction (qRT-PCR)

At 6, 24 and 48 h postreperfusion, the ischemic penumbra cortices were rapidly dissected on ice. Total RNA from the tissue samples was extracted using a TaKaRa MiniBEST Universal RNA Extraction Kit (TaKaRa, Japan) according to the manufacturer’s instructions. For cDNA synthesis, 1 μg total RNA was reverse transcribed into cDNA using the PrimeScriptTM RT reagent kit with gDNA Eraser (Perfect Real Time) (TaKaRa, Japan) according to the manufacturer’s instructions. Real-time monitoring of the PCR amplification reaction was performed using an ABI 7300 fast real time PCR system (CA, USA) and SYBR®Premix Ex TaqTM (Tli RNaseH Plus) (TaKaRa, Japan). The primers used were as follows: Fis1 mRNA (Forward: 5’-CTGGACTCATTGGACTGGCTGTG-3’; Reverse: 5’-AGGAAGGCGGTGGTGAGGATG-3’); β-actin (Forward:5’-CACCCGCGAGTACAACCTTC-3’; Reverse: 5’-CCCATACCCACCCATCACACC-3’). The relative quantitative value was determined using the 2^-ΔΔCt^ method, and β-actin expression was used as an internal control.

### Infarct volume percentage analysis

After 24 h of reperfusion, animals in each group were decapitated, and the brains were quickly removed, placed in a refrigerator at −20 °C for 20 min, sectioned into coronal slices of 2 mm thickness, and stained with 2,3,5-triphenyl-2H-tetrazolium chloride solution (TTC) (Amresco, Solong, CA, USA) at 37 °C for 30 min in the dark to evaluate the infarct area. Normal brain tissue was stained red, whereas infarcted tissue was stained pale gray. The sections were photographed and analyzed using ImageJ software (NIH, Maryland, USA). The cerebral infarct size was calculated as follows: (left hemisphere volume – right noninfarct volume)/left hemisphere volume×100%.

### Transmission electron microscopy

After 24 h of reperfusion, the brains were perfusion-fixed with 2.5% glutaraldehyde. Coronal brain sections (1×1×1 mm^3^) at the parietal lobe with cerebral ischemia were postfixed with 4% glutaraldehyde at 4 °C for 2 h. The sections were then rinsed in 0.2 M PBS 3 times, soaked in 1% osmium tetroxide for 2 h, rinsed in 0.2 M PBS again, dehydrated in an ascending ethanol series to 100%, and embedded in epoxy resin. The sections were sectioned into 50 nm ultrathin sections with an ultramicrotome (Leica, UC6, Wetzlar, Germany) and placed on 200-mesh copper grids. Then, the ultrathin sections were stained with lead citrate followed by observation under an H-7650 transmission electron microscope (Hitachi, Hiyoda, Tokyo).

### Statistical analysis

All data are expressed as the mean±standard deviation (*χ* − ±s) and analyzed using SPSS 19.0 statistical software (IBM Corporation, Armork, NY, USA). One-way analysis of variance (ANOVA) was used for comparison between different groups followed by LSD posttests. *P*<0.05 was considered statistically significant.

## Results

### Perfusion of cold saline via the carotid artery selectively reduced cortical temperature

We first explored whether cold saline perfusion via the carotid artery induced SBH. Therefore, we continuously monitored cortical and rectal temperatures for 1 h after reperfusion. In the hypothermia (HT) group, cortical temperature rapidly dropped (Fig 1A, from 34.39±0.23 °C to 32.22±0.09 °C, *P*<0.01) during the infusion period and was maintained below 33.0 °C for at least 15 min after the infusion ended. However, no significant changes in cortical temperature were found in the normothermia (NT) group (Fig 1A, from 34.55±0.30 °C to 34.81±0.44 °C, *P*>0.05). Rectal temperature did not change during the observational period in either the HT group or the NT group (Fig 1B, *P*>0.05). These results suggested that cold saline treatment achieved SBH.

**Fig 1.**
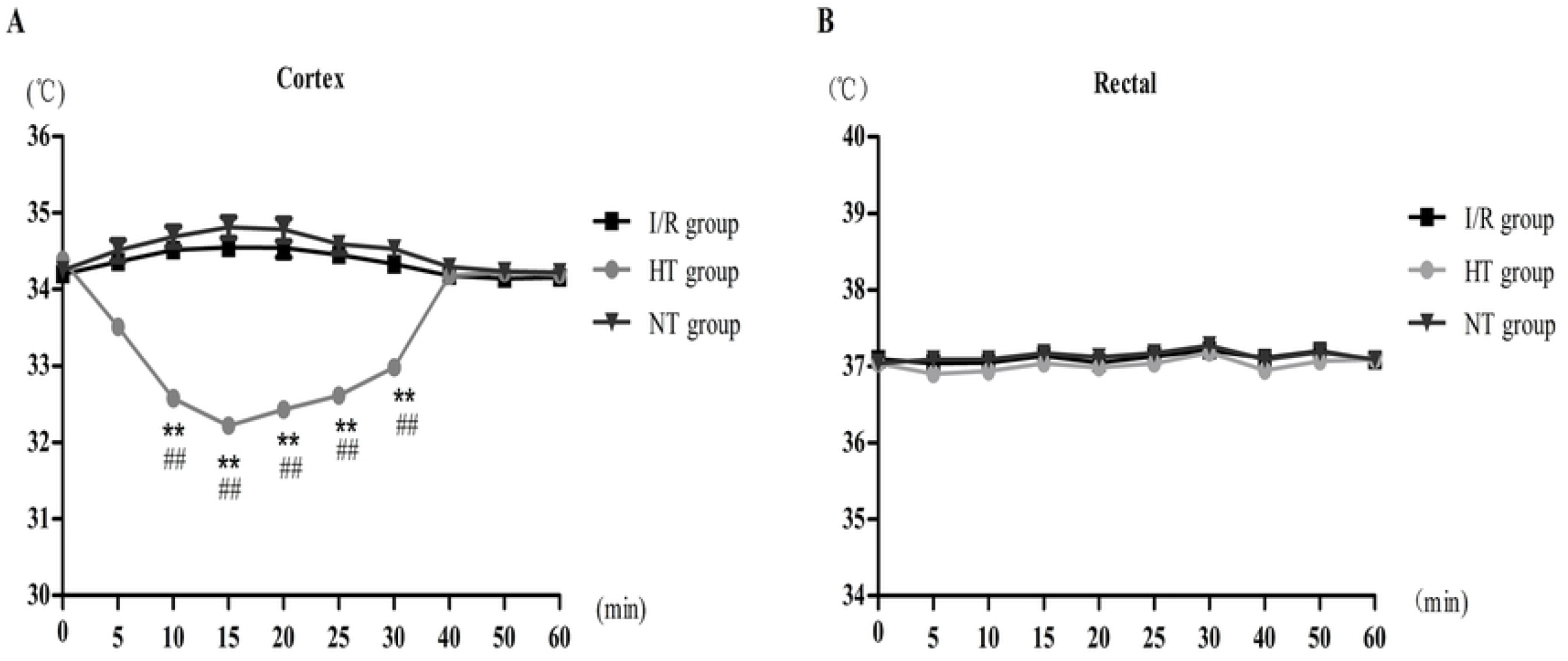
The cortical and rectal temperatures in rats with focal cerebral I/R injury 1 h after reperfusion. (A) Cortical temperature and (B) rectal temperature. One-way analysis of variance (ANOVA) was used to compare different groups followed by LSD posttests. Data are shown as the mean ± SEM. \*\**P* < 0.01 versus I/R group; ^##^*P* < 0.01 versus NT group. I/R: ischemia/reperfusion; HT: hypothermia; NT: normothermia, n = 10 each.

### SBH decreased neurological deficit scores (NDS) and cerebral infarct size

In this study, the rat neurological examination was conducted using the Zea Longa 5-point scoring method at 6, 24 and 48 h postreperfusion (Fig 2). The cerebral ischemia infarct size was determined by 2,3,5-triphenyl-2H-tetrazolium chloride solution (TTC) staining at 24 h after reperfusion (Fig 3A). Higher NDS and cerebral infarct size were observed in the I/R, HT and NT groups than in the sham group (*P*<0.05). After the cold saline treatment in the HT group, NDS and cerebral infarct size decreased (*P*<0.05). Interestingly, there was hardly any difference in NDS and cerebral infarct size between the NT group and I/R group (*P*>0.05) (Figs 2 and 3B).

**Fig 2.**
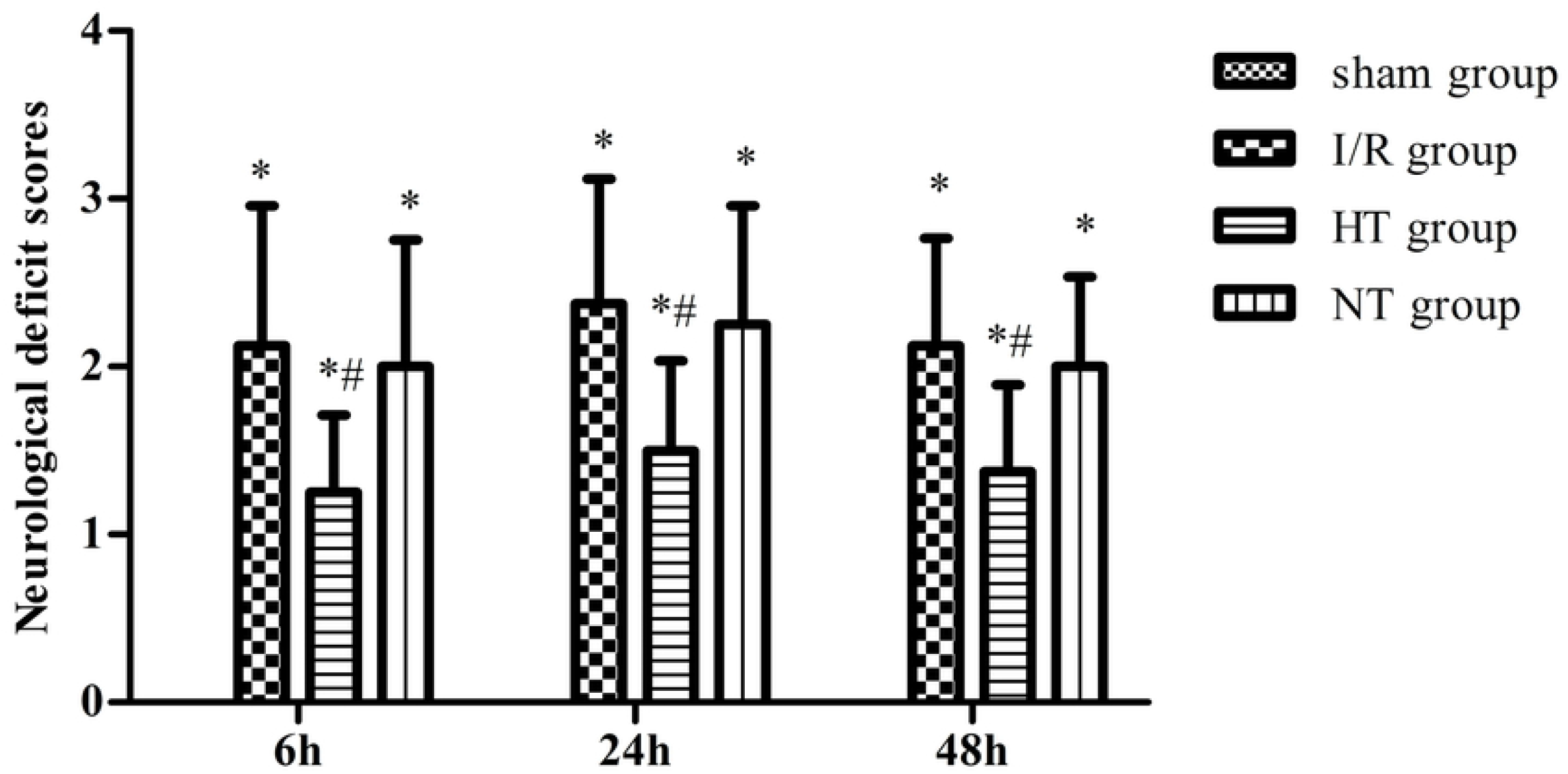
Effects of SBH on NDS in rats with focal cerebral I/R injury. One-way analysis of variance (ANOVA) was used to compare different groups followed by LSD posttests. Data are shown as the mean ± SD. \**P* < 0.05 versus sham group; #*P* < 0.05 versus I/R group. I/R: ischemia/reperfusion; HT: hypothermia; NT: normothermia, n = 8 each.

**Fig 3.**
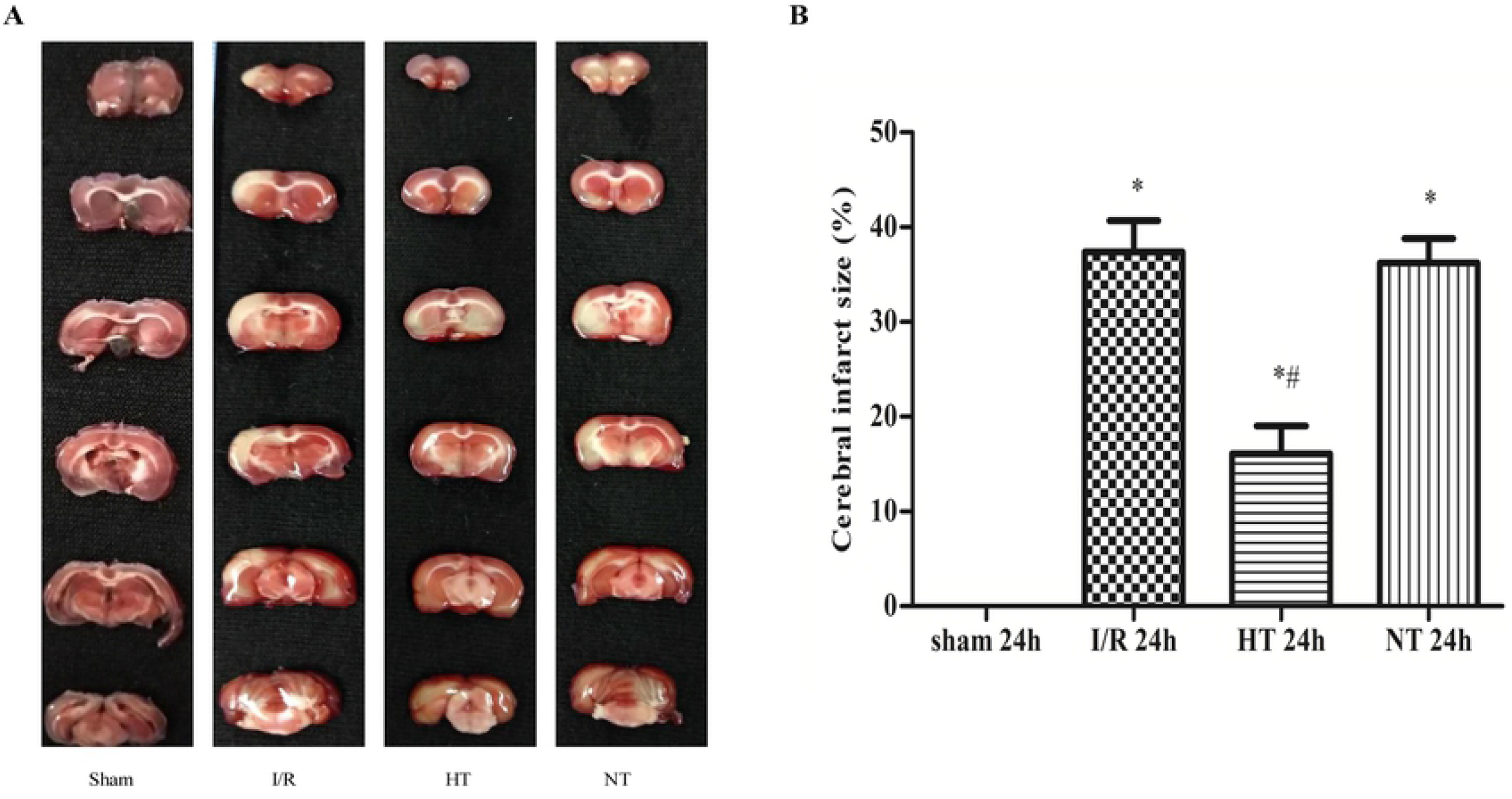
Effects of SBH on infarct volume in rats with focal cerebral I/R injury. (A) Representative TTC-stained coronal brain sections. (B) Quantification of the percentage of cerebral infarct size among the four groups. One-way analysis of variance (ANOVA) was used to compare different groups followed by LSD posttests. Data are shown as the mean ± SD. \**P* < 0.05 versus sham group; #*P* < 0.05 versus I/R group. TTC: 2,3,5-triphenyl-2H-tetrazolium chloride solution; I/R: ischemia/reperfusion; HT: hypothermia; NT: normothermia, n = 5 each.

### SBH alleviated histopathological changes and neuronal apoptosis in the cortical penumbra caused by I/R

At 6, 24 and 48 h postreperfusion, hematoxylin-eosin (HE) staining showed that no abnormally morphological cells were detected in the sham group, while the other three groups showed shrunken cell bodies and nuclear pyknosis. By contrast, SBH markedly ameliorated pathological changes in the HT group compared with those in the I/R and NT groups (Fig 4A). Terminal deoxynucleotidyl transferase-mediated dUTP nick-end labeling (TUNEL) analysis was used to detect neuronal apoptosis. As shown in Figs 4B and 4C, the proportion of TUNEL-positive cells showed an increase in neuronal apoptosis in the ischemic penumbra at 6, 24 and 48 h postreperfusion in the three injury groups compared to that in the sham group (*P*<0.05). Conversely, the proportion of TUNEL-positive cells in the HT group was lower than that in the I/R group at 6, 24 and 48 h postreperfusion (*P*<0.05). However, there were no obvious changes between the NT group and I/R group (*P*>0.05).

**Fig 4.**
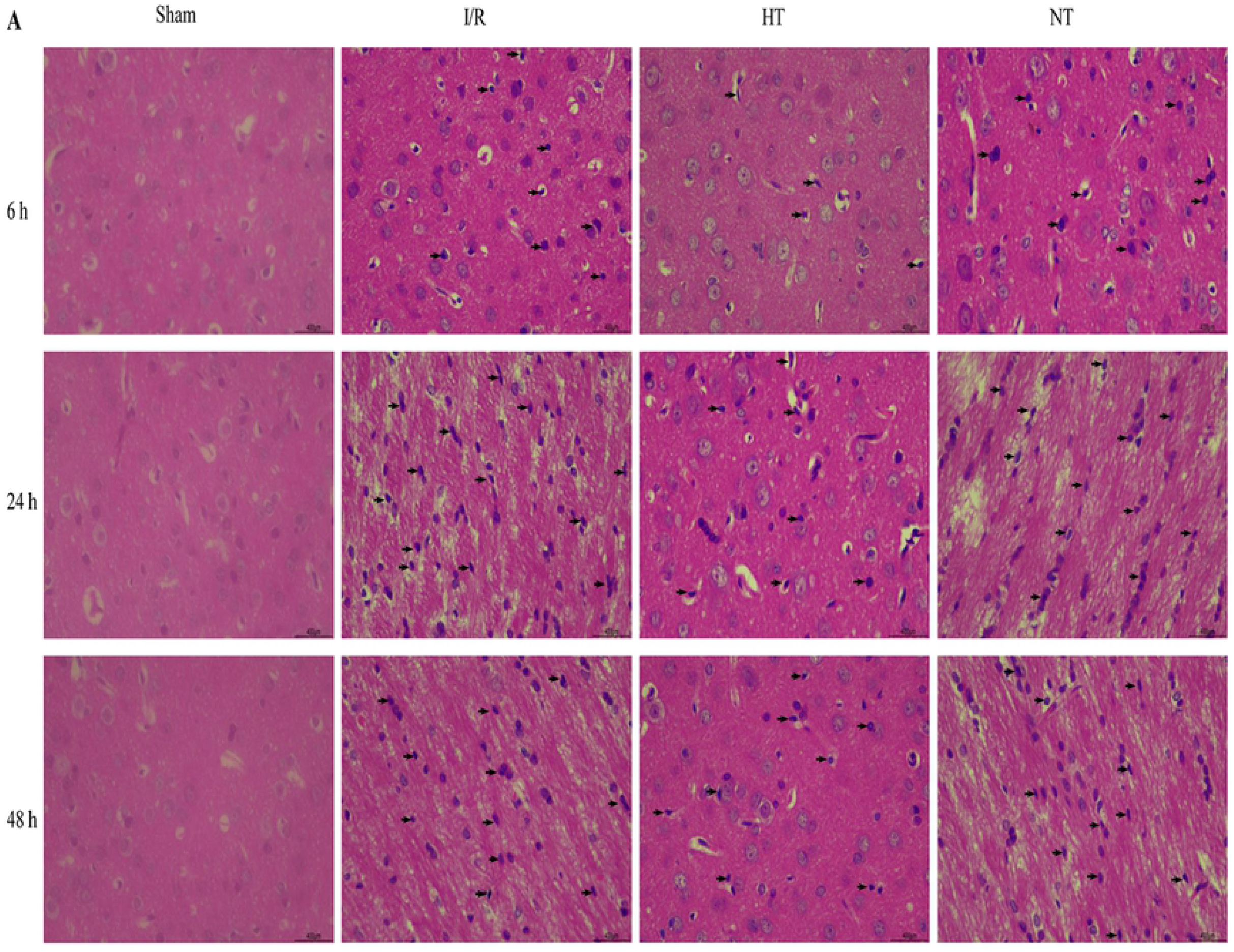
Effects of SBH on ischemia-induced histopathological changes and neuronal apoptosis. (A) Representative pictures of HE staining in the cortical ischemic penumbra. Shrunken cell bodies and nuclear pyknosis (black arrows) were observed in the I/R, HT and NT groups but not in the sham group. The degree of histopathological changes in the HT group was slight less than that in the I/R group. (B) Representative pictures of TUNEL staining showed neuronal apoptosis in the cortical ischemic penumbra. TUNEL-positive nuclei with brownish yellow granules were considered apoptotic cells as indicated by black arrows. (C) Quantification of the percentage of TUNEL-positive cells among the four groups. Data are expressed as the mean ± SD. \**P* < 0.05 versus sham group; #*P* < 0.05 versus I/R group. Scale bars = 400 μm, 400× visual field. I/R: ischemia/reperfusion; HT: hypothermia; NT: normothermia, n = 5 each.

### SBH decreased the expression of Fis1 protein and mRNA in the ischemic penumbra caused by I/R

The expression of Fis1 in the ischemic penumbra was measured by Western blot analysis, and the expression of Fis1 mRNA was measured by qRT-PCR. Parallel variation trends in the expression of Fis1 protein and mRNA were observed. The expression of Fis1 protein and mRNA was higher at 6, 24 and 48 h postreperfusion in the three injury groups than in the sham group (P<0.05). The expression of Fis1 protein and mRNA at 6, 24 and 48 h postreperfusion declined following the infusion of cold saline in the HT group (P<0.05). However, there were hardly any changes in the expression of Fis1 protein and mRNA between the NT group and I/R group (*P*>0.05) (Figs 5B and 5C).

**Fig 5.**
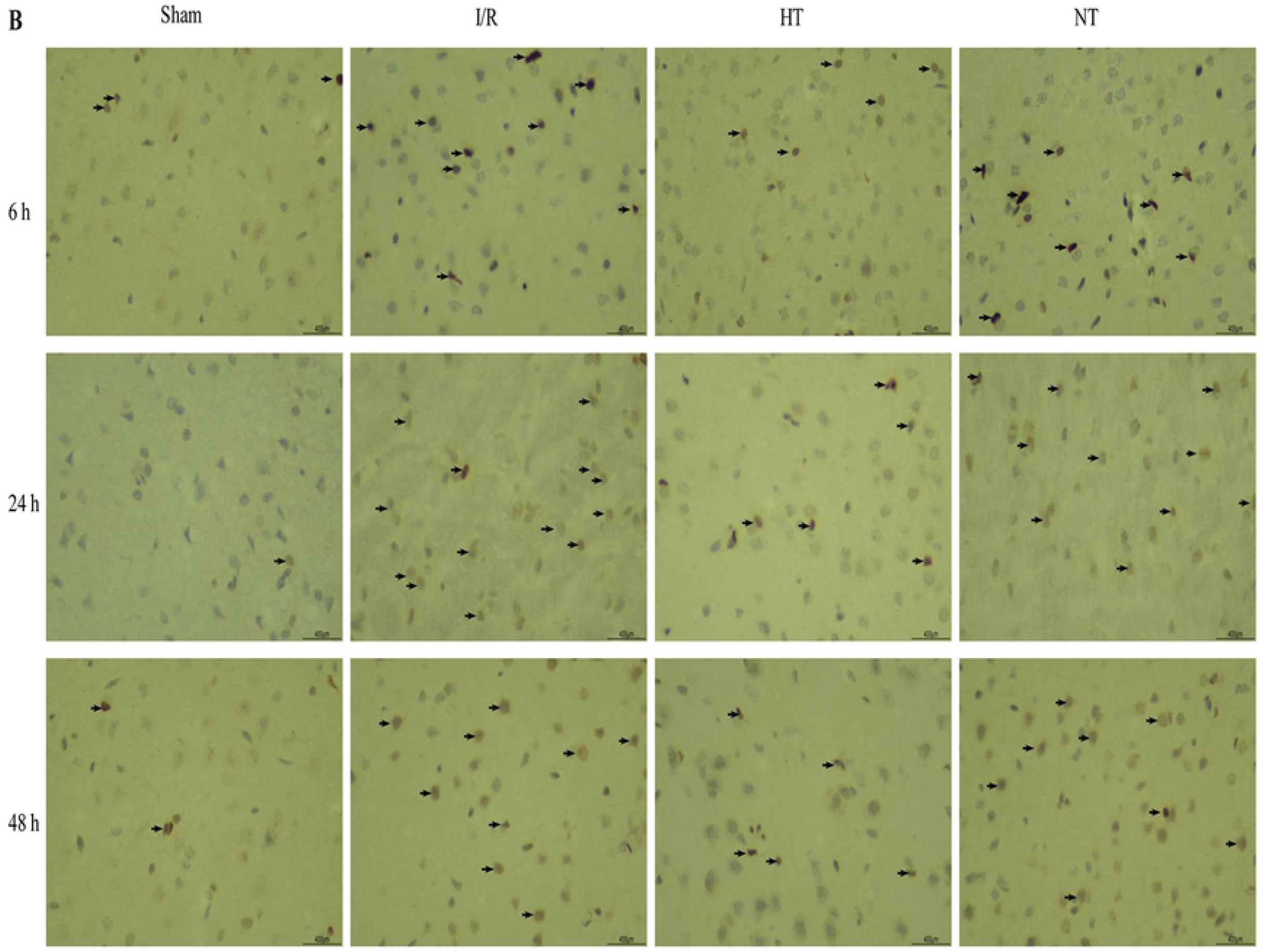
Expression of Fis1 in the cortical ischemic penumbra following focal cerebral I/R injury in rats. (A) Western blotting was used to measure Fis1 protein from the cortical ischemic penumbra. Blots are representative of rats per group. (B) The relative quantities of Fis1 to β-actin. The results were normalized to the percentage of β-actin expression. (C) The expression of Fis1 mRNA from the cortical ischemic penumbra was determined by qRT-PCR. The relative quantities of Fis1 mRNA to β-actin. The results were normalized to the percentage of β-actin expression. One-way analysis of variance (ANOVA) was used to compare different groups followed by LSD posttests. Data are shown as the mean ± SD. \**P* < 0.05 versus sham group; #*P* < 0.05 versus I/R group. I/R: ischemia/reperfusion; HT: hypothermia; NT: normothermia, n = 5 each.

### SBH reduced cyto-Cyto c expression in the ischemic penumbra caused by I/R

Cyto c is released from mitochondria to the cytosol during mitochondrial fission, which can induce neuronal apoptosis. Therefore, we analyzed the expression of cyto-Cyto c by Western blotting to determine the level of mitochondrial fission. As shown in Fig 6B, the expression of cyto-Cyto c was significantly increased at 6, 24 and 48 h postreperfusion in the three injury groups compared to that in the sham group (*P*<0.05). By contrast, the expression of cyto-Cyto c at 6, 24 and 48 h postreperfusion declined following the infusion of cold saline in the HT group (*P*<0.05). However, no significant changes in the expression of cyto-Cyto c were found between the NT group and I/R group (*P*>0.05).

**Fig 6.**
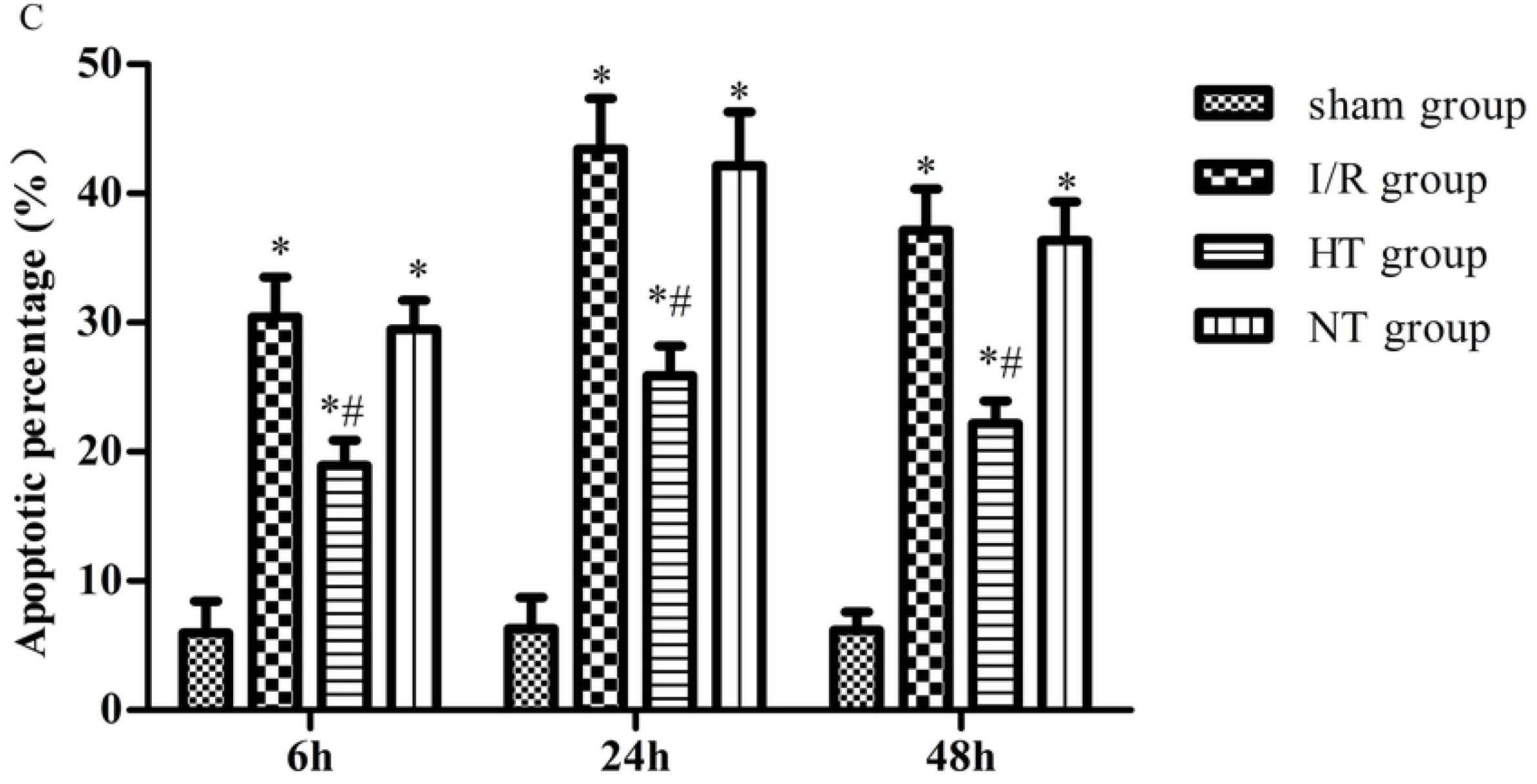
Expression of cyto-Cyto c in the cortical ischemic penumbra following focal cerebral I/R injury in rats. (A) Western blotting was used to measure cyto-Cyto c from the cortical ischemic penumbra. Blots are representative of rats per group. (B) The relative quantities of cyto-Cyto c to β-actin. The results were normalized to the percentage of β-actin expression. One-way analysis of variance (ANOVA) was used for comparison between different groups followed by LSD posttests. Data are shown as the mean ± SD. \**P* < 0.05 versus sham group; #*P* < 0.05 versus I/R group. I/R: ischemia/reperfusion; HT: hypothermia; NT: normothermia, n = 5 each.

### Mitochondria ultrastructural alterations

Transmission electron microscopy was used to detect the mitochondrial ultrastructure of neurons at 24 h after reperfusion. The mitochondria were well arranged and exhibited a complete bilayer membrane structure and normal cristae without swelling and vacuolar degeneration in the sham group (Fig 7A). By contrast, mitochondrial morphology showed obvious alterations such as disappearance of the bilayer membrane structure, vacuolar degeneration and swelling and loss of cristae. These outcomes indicated that excessive mitochondrial fission occurred at 24 h after reperfusion in the I/R and NT groups (Figs 7B and 7D). However, these detrimental morphological changes were ameliorated, and mitochondria exhibited a less swollen and relatively intact membrane in the HT group (Fig 7C).

**Fig 7.**
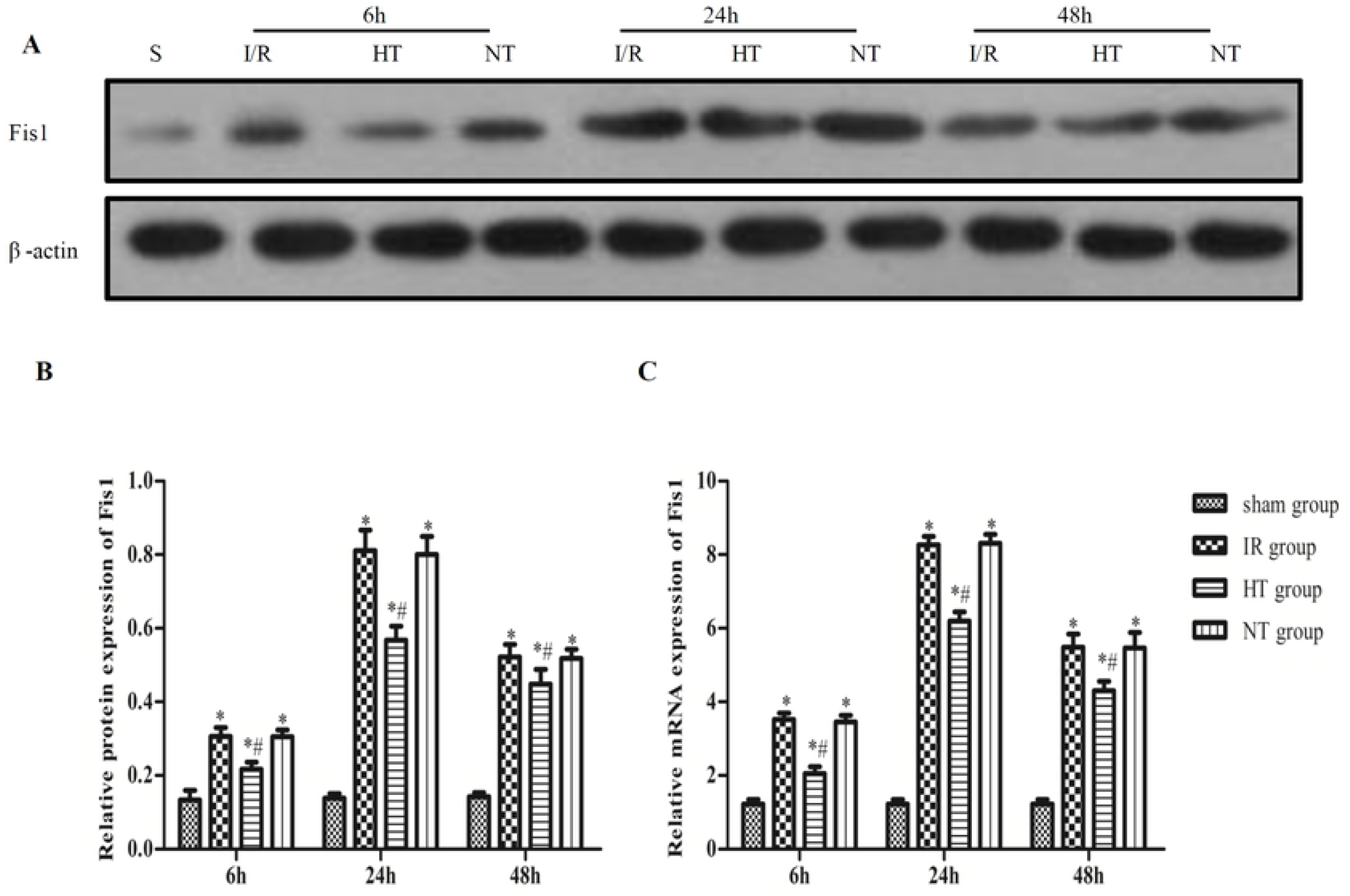
Representative electron photomicrographs for mitochondrial shape and ultrastructure (black arrow) in the cortical ischemic penumbra. (A) shows many profiles of normal mitochondria in the sham group. (B) and (D) show a large number of mitochondria with swelling and disorganized cristae, as well as partial vacuolar degeneration in mitochondria in the I/R and NT groups. (C) shows slight mitochondrial swelling and disorganized cristae, which reflects mild mitochondrial damage after cold saline infusion in the HT group. Scale bars = 1 μm. I/R: ischemia/reperfusion; HT: hypothermia; NT: normothermia, n = 5 each.

## Discussion

Cerebral I/R injury is a common clinical pathophysiological phenomenon after restoring blood perfusion in stroke patients, which involves multiple pathogenesis [7]. Mitochondria reportedly play an important role in the development of I/R injury, which is involved in calcium homeostasis, oxidative phosphorylation, reactive oxygen species production, and apoptosis [13]. Recent studies have suggested that morphological changes in the mitochondria are relevant to I/R injury [8,22,23]. In stress, the balance of mitochondrial fission and fusion is lost, and fission prevails, which promotes fragmented mitochondria and induces mitochondrial dysfunction [24–26]. Moreover, in recent reports, excessive mitochondrial fission or fragmentation was observed during cerebral I/R injury [8,27,28]. In addition, Fis1, as one of the mitochondrial fission systems, plays an important role in the regulation of mitochondrial fission. Fis1 is exclusively localized to the mitochondrial outer membrane [14] and mediates mitochondrial fission by recruiting cytoplasmic Drp1. In our previous study, the expression of Drp1 in the mitochondrial outer membrane increased during cerebral I/R injury [21]. Based on this finding, we was speculated that the expression of Fis1 would increase accordingly during cerebral I/R injury. We examined Fis1 protein and mRNA in our study to test this hypothesis. Interestingly, some reports also confirmed our hypothesis by demonstrating that Fis1 expression increased following cerebral I/R injury [8,29].

Apoptosis is considered a vital component of the development of cerebral I/R injury [30]. Excessive mitochondrial fission promotes mitochondrial outer membrane permeability and increases Cyto c release, subsequently activating the apoptotic cascade reaction and eventually aggravating neurological damage [13,31]. Moreover, in a study by Wang et al. [32], adenoviral Fis1, which can induce Fis1 overexpression, increased mitochondrial fission and apoptosis, whereas Fis1 knockdown attenuated mitochondrial fission and apoptosis. In addition, mitochondrial morphological alterations, cyto-Cyto c expression and neuronal apoptosis were observed in our experiment. Consistent with previous reports, our results further demonstrated that Fis1 overexpression increased mitochondrial fission, causing Cyto c release and apoptosis during cerebral I/R injury.

Notably, many studies have focused on the neuroprotection of hypothermia, one of the most robust neuroprotectants against ischemia stroke [33–36]. SBH is more suitable for neuroprotection after stroke because it can quickly reach the target temperature and avoid the adverse effects associated with general hypothermia [37,38]. Moreover, transarterial regional hypothermia, which can achieve SBH, may exert strong neuroprotective effects in the MCAO with transient collateral hypoperfusion model [19]. In the present study, we prepared a hypothermia model following MCAO by perfusing cold saline through the ICA, and the results of cortical and rectal temperatures showed that SBH models were successfully established. To eliminate interference from hemodilution by saline infusion, we performed 37 °C warm saline infusion in the same manner in the NT group. Our data demonstrated that the NDS and cerebral infarct volume percentage decreased after cold saline perfusion in the HT group. By contrast, there were hardly any differences in the results between the I/R group and NT group. These results showed that SBH can attenuate focal cerebral I/R injury.

However, the underlying mechanisms of hypothermia-induced neuroprotection are complex and remain unclear. Notably, morphological changes in the mitochondria are a key step of cerebral I/R injury, and mitochondria are a target of hypothermia-induced neuroprotection. General hypothermia can attenuate mitochondrial oxidative stress and reduce mitochondrial membrane permeability [39,40]. In addition, in our previous study, general hypothermia reduced mitochondrial fission by inhibiting the translation of Drp1 from the cytoplasm to the mitochondrial outer membrane [17]. Drp1 is mainly localized to the cytoplasm, and mitochondrial fission is inhibited in cells with Fis1 deletion mutations [16]. However, whether SBH could inhibit mitochondrial fission and subsequently decrease neuronal apoptosis by inhibiting the expression of Fis1 has not been thoroughly studied. Therefore, in our study, we analyzed the expression levels of Fis1 protein and mRNA and cyto-Cyto c, as well as the ratio of neuronal apoptosis in the ischemic penumbra of the cerebral cortex in rats with focal cerebral I/R injury. Interestingly, the results clearly demonstrated that SBH inhibited the expression of Fis1 protein and mRNA and cyto-Cyto c, in addition to the ratio of neuronal apoptosis at 6, 24 and 48 h after reperfusion. Furthermore, mitochondrial ultrastructural analysis revealed that at 24 h after reperfusion, the level of mitochondrial fission was lower in the HT group than in the I/R group and NT group. Combined with our present research and a previous report [17], we hypothesized that SBH could down-regulate the expression of Fis1 in the mitochondrial outer membrane, inhibiting excessive mitochondrial fission induced by the binding of Drp1 to Fis1, reducing the cytosolic release of Cyto c and eventually ameliorating cellular apoptosis.

Although the neuroprotective mechanisms of SBH against I/R injury have not been fully examined in this work, our study demonstrated that SBH might play a neuroprotective role by regulating the expression of Fis1 to some extent. However, this study has some limitations. First, the precise mechanisms of SBH-induced neuroprotection by suppressing Fis1 remain unclear. We will implement Fis1 overexpression and knockdown in vitro and vivo in our future study. Second, further studies are needed to investigate the changes in the degree of binding of Drp1 to Fis1 after SBH.

In conclusion, SBH could ameliorate focal cerebral I/R injury through inhibiting Fis1 expression and mitochondrial fission.

## Acknowledgments

We would like to thank the teachers and students at Qingdao Municipal Hospital for their assistance.

## Author Contributions

**Data curation:** Yanan Tang.

**Formal analysis:** Yanan Tang, Gaofeng Zhang.

**Investigation:** Yanan Tang, Gaofeng Zhang, Mingshan Wang.

**Methodology:** Yanan Tang, Gaofeng Zhang, Mingshan Wang

**Project administration:** Yanan Tang, Weiwei Qin, Huailong Chen.

**Resources:** Yanan Tang, Weiwei Qin, Xiaopeng Sun, Fei Shi, Lixin Sun.

**Software:** Yanan Tang, Xiaona Xu.

**Supervision:** Yanan Tang, Gaofeng Zhang, Mingshan Wang.

**Validation:** Yanan Tang, Gaofeng Zhang, Mingshan Wang.

**Writing – original draft:** Yanan Tang

**Writing – review & editing:** Yanan Tang, Gaofeng Zhang, Mingshan Wang.

## Supporting information

**S1 Methods.**
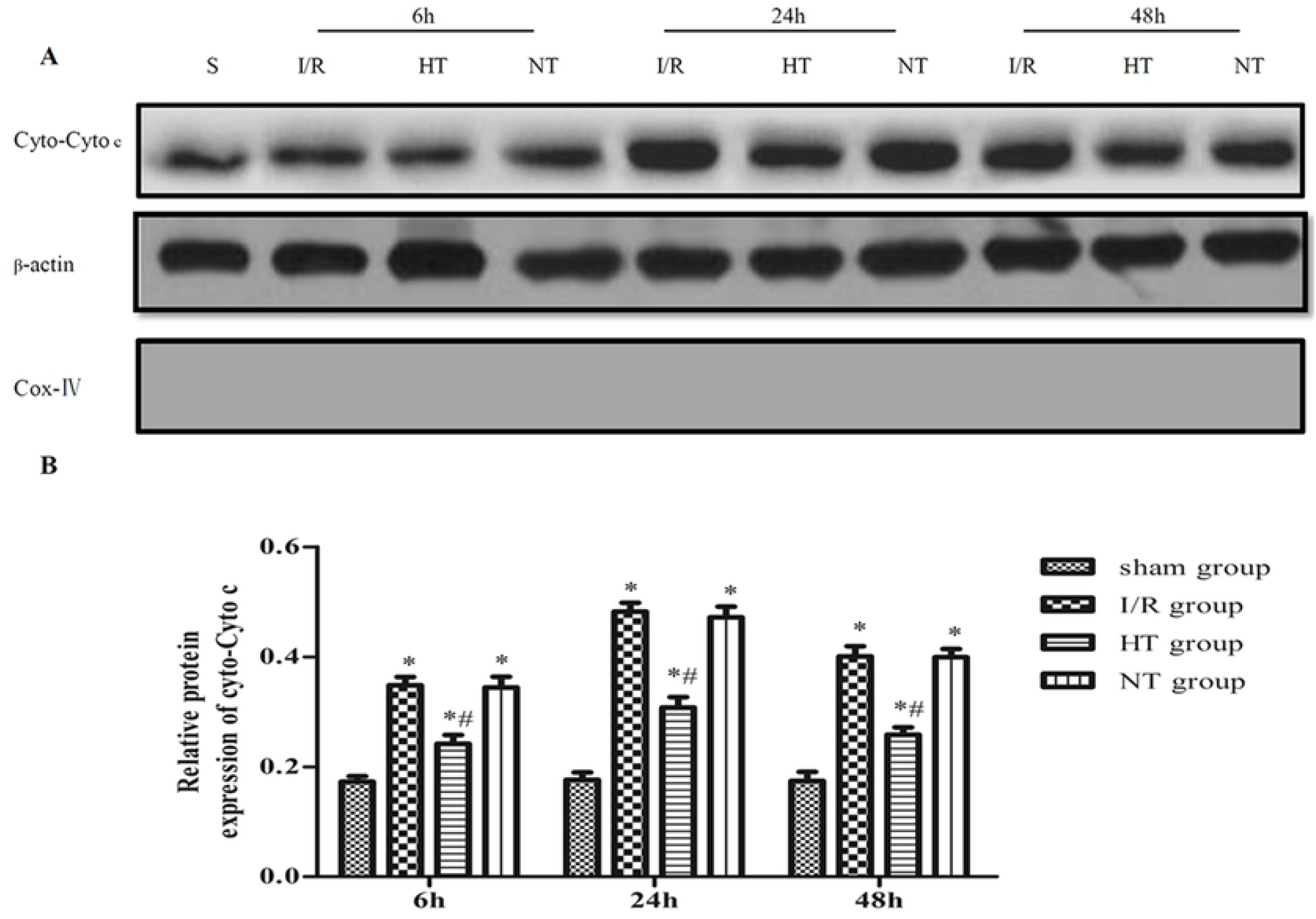
Establishment of the focal cerebral I/R injury model. (DOCX)

**S2 Methods.**
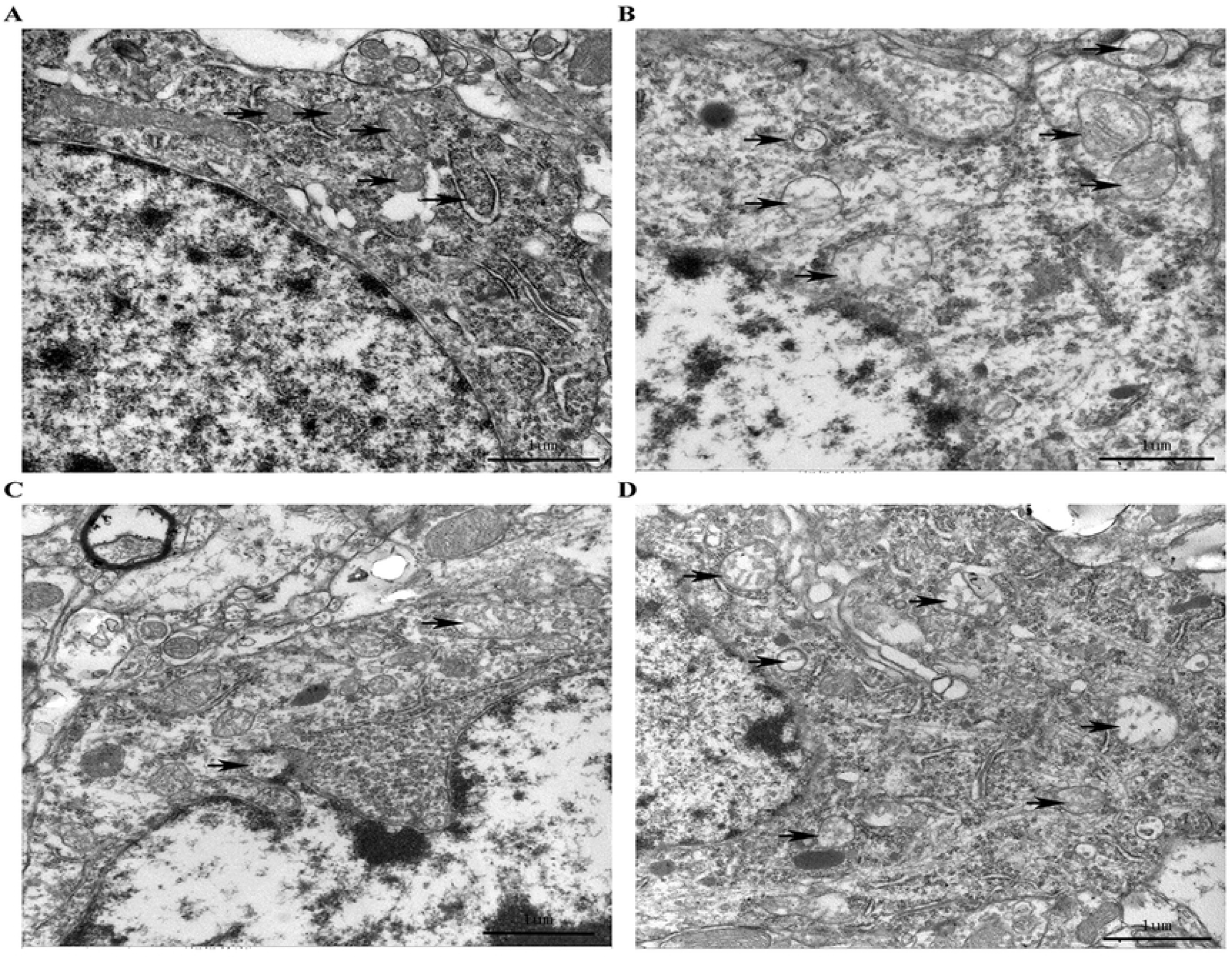
Steps for isolating cortical ischemic penumbra. (DOCX)

